# Synergism between potassium and nitrate enhances the alleviation of ammonium toxicity in rice seedling root

**DOI:** 10.1101/2021.03.08.434377

**Authors:** Gen Fang, Jing Yang, Tong Sun, Xiaoxin Wang, Yangsheng Li

**Affiliations:** State Key Laboratory of Hybrid Rice, College of Life Sciences, Wuhan University, Wuhan, China

**Author notes:** Corresponding author, (YL).

**Keywords:** Rice, Root, Synergism, Ammonium toxicity alleviation, Potassium, Nitrate

## Abstract

Ammonium toxicity in plants is considered a global phenomenon, but the primary mechanisms remain poorly characterized. Here, we showed that although the addition of potassium (K^+^) or nitrate (NO_3_^−^) partially alleviated the inhibition of rice root growth caused by ammonium toxicity, the coexistence of K^+^ and NO_3_^−^ clearly improved the alleviation via a synergistic mechanism. The synergism led to significantly improved alleviation effects on root biomass, length, surface area, number and meristem cell number. The aberrant auxin distribution in root tips, rhizosphere acidification level and abnormal cell morphology in the root cap and elongation zone caused by ammonium toxicity could be recovered by this synergism. RNA sequencing and the weighted gene correlation network analysis (WGCNA) revealed that the mechanism of this synergism involves cellulose synthesis, auxin and gibberellin metabolism regulation at the transcription level.

## Introduction

Compared with nitrate, ammonium is the preferred nitrogen source for paddy rice and many other plant species. While low supply of ammonium promotes plant growth, high supply of ammonium causes toxicity, especially when applied alone [1]. Ammonium toxicity can lead to plant growth retardation, biomass reduction, structural changes, the formation of short and dense root systems and even death [2]. In addition, ammonium toxicity inhibits root cell elongation and affects root geotropism by disturbing auxin distribution in root tips. [3–5]. Cells that absorb excess ammonium actively excrete the ammonium. These transport processes take up a large number of ion channels, resulting in a large waste of energy [6]. To maintain charge balance, plant cells need to pump out protons to counteract the excess ammonium in the cytoplasm, which causes rhizosphere acidification [7]. Ammonium can enter cells through a variety of nonselective cation channels and potassium-specific channels, which competitively inhibit the uptake of potassium [8, 9]. Excessive ammonium assimilation in the cytoplasm consumes a large amount of carbonic compounds, which results in a lack of carbon sources in plants [10]. The auxin signal in Arabidopsis roots was obviously reduced under ammonium treatment, and exogenous application of auxin could not completely alleviate the toxicity phenotype in the roots [11]. Ammonium toxicity can result in higher lipid peroxidation levels, a higher ratio of the oxidized state of glutathione and ascorbic acid, and higher activity of ROS-scavenging enzymes [12].

The way to alleviate plant ammonium toxicity is to reverse these toxic mechanisms, for instance, by increasing the activity of the ammonium assimilation system, relieving rhizosphere acidification, inhibiting the futile transmembrane ammonium cycling, increasing the supply of other nutrients [1, 2, 13]. Potassium has a significant alleviation effect on ammonium toxicity [4]. A possible reason for this effect is that potassium can competitively suppress the transport of ammonium, thereby reducing futile transmembrane ammonium cycling [14]. Potassium is important to protein anabolism. It affects the synthesis and signaling of auxin, similar to ammonium. Some researchers have reported that the presence of nitrate in the growth medium restores the ammonium toxicity phenotype in the plant, even at a very low nitrate concentration [8]. Since nitrate is an essential signaling molecule for plants, the alleviation effect of nitrate on ammonium toxicity could be extremely complex [15]. Nitrate, as an anionic and oxidized nitrogen source, can mitigate the imbalance of ions and redox states caused by cations and reduce ammonium ions when applied together [16]. The rhizosphere becomes alkalized when plants absorb nitrate, which counteracts the acidification caused by ammonium absorption [13]. Nitrate alleviates ammonium toxicity without lessening ammonium accumulation, organic acid depletion and inorganic cation depletion in *Arabidopsis thaliana* shoots [17].

In this study, we compared the alleviation effects of K^+^/NO_3_^−^/KNO_3_ on ammonium toxicity, and found that combined application of K^+^ and NO_3_^−^ not only showed improved alleviation of ammonium toxicity compared to application of individual ions, but also resulted in some novel beneficial mechanisms, probably due to an undiscovered synergism. We provide morphological and physiological evidence of the existence of this synergism, and we used transcriptome analysis to reveal a regulatory network for cell wall construction related to this synergism.

## Materials and Methods

### Plant materials and growth conditions

The plant materials used in this research included ZH11 (*Oryza sativa japonica*) and DR5:GUS insertion lines (ZH11 background).

Rice seeds were germinated in distilled water for 2 days in a dark environment at 30°C and then transferred into different hydroponic culture media in 400-mL boxes (24 plants per box) and grown in a growth chamber (30°C-14 hour-light/22°C-8 hour-dark cycles, 100 μmol photons m^−2^ s^−1^ illumination and 80% humidity).

There were 6 groups of hydroponic culture media used in this research: control (pure water); ammonium toxicity treatment (4 mM NH_4_Cl solution); K^+^ alleviation treatment (4 mM NH_4_Cl and 4 mM KCl solution); NO_3_^−^ alleviation treatment (4 mM NH_4_Cl and 4 mM NaNO_3_ solution); K^+^ and NO_3_^−^ synergistic alleviation treatment (4 mM NH_4_Cl and 4 mM KNO_3_ solution); Na^+^ and Cl^−^ alleviation treatment (4 mM NH_4_Cl and 4 mM NaCl solution). No other nutrients existed in the hydroponic culture media except for the ions mentioned above.

If not mentioned, 7-day-old seedlings were harvested for follow-up experiments.

### Analysis of root morphology

The whole root area per plant was photographed by a D7000 camera (Nikon, Japan). All the root morphology parameters were measured by SmartRoot software V4.2 [18] according to its user guide.

The root tips were cut and fixed with FAA fixative (1.9% formaldehyde, 4.9% acetic acid, 63% ethanol, m/V) for 2 days and then embedded in paraffin. Then, 6-μm-thick slices were obtained with a microtome. After dewaxing with xylene, the sections were placed on glass slides and photographed using a microscope (DM4000B equipped with a DFC490 camera, Leica, Germany).

### Culture pH measurement

On day 0/3/5/7 after germination, the pH of the hydroponic culture media of the 5 treatments was measured using a pH meter (PB-10, Sartorius, Germany).

### DR5:GUS-based auxin distribution assay

DR5:GUS insertion lines grown under the 5 different treatments were harvested and rinsed in distilled water for 30 s. After fixing in 90% (m/m) acetone in vacuo, the roots were rinsed in GUS staining buffer (50 mM Na_2_HPO_4_, 50 mM NaH_2_PO_4_, 0.1% Triton X-100, 2 mM K_4_[Fe(CN)_6_], 10 mM EDTA) 3 times, and then, they were stained in 5 mL of GUS staining buffer containing 5 μg of X-gluc (Sigma-Aldrich, USA). Images were photographed by a microscope (DM4000B equipped with a DFC490 camera, Leica, Germany).

### Auxin alleviation treatment

After germination, rice seeds were transferred into 4 mM NH_4_Cl containing 0/10/15/20 nM IAA/NAA and grown for 14 days. The radicle lengths were measured on day 14.

### Ion content, enzyme activity and soluble protein content determination in root tissues

Seedling roots were harvested and rinsed in distilled water for 30 s to remove extracellular ions. Roots were homogenized under liquid N_2_ using a mortar and pestle.

For tissue NH_4_^+^ content determination, 0.2 g of homogenized tissue was mixed with 2 mL of 10 mM precooled formic acid before centrifugation (4°C, 10000 ×g, 10 minutes). The supernatants were analyzed by the o-phthalaldehyde (OPA) method to determine the NH_4_^+^ content using spectrophotometry as described in [19].

For tissue NO_3_^−^ content determination, 0.1 g of homogenized tissue was mixed with 1 mL H_2_O before centrifugation (4°C, 10000 ×g, 10 minutes). Then, 30 μL of supernatant was mixed with 100 μL of 5% (m/v) salicylic acid-H_2_SO_4_. After reacting for 20 minutes at room temperature, 2 mL of 2 M NaOH was added to the mixture. Sample absorbance was measured at 410 nm, and the NO_3_^−^ content was computed according to a standard curve.

The K^+^ content was determined by the atomic absorption spectroscopy (AAS) method using a contraAA700 (Analytik Jena, Thuringia, Germany). After determining the dry root weights, samples were digested with 5 mL of concentrated nitric acid and 1 mL of H_2_O_2_ at 150°C until the mixtures were transparent. Then, the solutions were diluted to 50 mL with 0.5% nitric acid. The K^+^ content was determined at 404.4 nm.

Homogenized tissue was lysed with Western and IP lysis buffer (Beyotime Biotechnology, Shanghai, China). The soluble protein content was measured with a BCA protein assay kit (Beyotime Biotechnology, Shanghai, China).

GS activity was measured using a GS activity assay kit (Comin Biotechnology, Suzhou, China) as described in [20].

GOGAT activity was measured using a GOGAT activity assay kit (Comin Biotechnology, Suzhou, China) as described in [21].

### Net ^15^NH_4_ flux measurement

Rice seedlings were germinated and grown on 4 mM ^14^NH_4_Cl for 7 days. After harvesting, their roots were rinsed with 0.1 mM CaSO_4_ immediately for 1 minute. Then, the samples were separated into 4 groups evenly and immersed in 4 mM ^15^NH_4_Cl, 4 mM ^15^NH_4_Cl + 4 mM KCl, 4 mM ^15^NH_4_Cl + 4 mM Na^14^NO_3_, or 4 mM ^15^NH_4_Cl + 4 mM K^14^NO_3_ for 1 hour. After rinsing again with 0.1 mM CaSO_4_ for 1 minute, the roots from the 4 groups were dried in a 70°C oven for 48 hours. Then, they were homogenized using liquid N_2_. The proportion of ^15^N in the total nitrogen content (m/m), which represents the root surface ^15^NH_4_ influx in 1 hour, was measured by isotope mass spectrometry (isoprime-100, Vario ISOTOPE cube CN).

Rice seedlings were germinated and grown on 4 mM ^15^NH_4_Cl solution for 7 days. After harvesting, their roots were rinsed with 0.1 mM CaSO_4_ immediately for 1 minute. Then, the samples were separated into 4 groups evenly and immersed in 4 mM ^14^NH_4_Cl, 4 mM ^14^NH_4_Cl + 4 mM KCl, 4 mM ^14^NH_4_Cl + 4 mM Na^14^NO_3_, or 4 mM ^14^NH_4_Cl + 4 mM K^14^NO_3_ for 1 hour. After rinsing again with 0.1 mM CaSO_4_ for 1 minute, the roots from the 4 groups were dried in a 70°C oven for 48 hours. Then, they were homogenized using liquid N_2_. The proportion of ^15^N in the total nitrogen content (m/m), which represent the root surface ^15^NH_4_ outflux in 1 hour, was measured by isotope mass spectrometry (isoprime-100, Vario ISOTOPE cube CN).

### Calculation of correlations between traits

Correlations between traits were calculated using the R package corrplot (order = “FPC”).

### RNA sequencing and data analysis

Total RNA was extracted from 15 sample of ZH11 roots under 5 different treatments (3 biological replications for each treatment, 40 plants for each biological replication). The cDNA libraries were constructed by Personal Biotechnology (Shanghai, China) using the TruSeq RNA Sample Prep Kit (Illumina, USA). mRNA paired-end sequencing was performed using the NextSeq 500 platform (Illumina, USA) by the same company. The raw RNA sequencing reads have been deposited into the NCBI Sequence Read Archive (SRA) under accession NO. PRJNA693667.

After removing the adapter sequences from raw reads, clean reads were obtained by Trimmomatic [22]. The clean reads were mapped to the reference genome of the *Oryza sativa* japonica variety (ftp://ftp.ensemblgenomes.org/pub/plants/release-36/fasta/oryza_sativa/dna/Oryza_sativa.IRGSP-1.0.dna.toplevel.fa.gz) using STAR [23], and gene structure information was obtained from EnsemblPlants annotation (ftp://ftp.ensemblgenomes.org/pub/plants/release-36/gtf/oryza_sativa/Oryza_sativa.IRGSP-1.0.36.chr.gtf.gz). The normalized gene expression levels (TMM normalized FPKM) were calculated using Trinity [24]. DESeq2 [25] was used to identify the DEGs. Those with an adjusted p-value (padj) < 0.05 and |log_2_FC| > 1 genes between every pair of groups were considered significant DEGs.

### WGCNA analysis

A matrix containing the TMM values of all genes was used as an input file, and 40% of the genes were first filtered out using the varFilter command (var.func = IQR) in the R package genefilter. Then, WGCNA was performed using the R package WGCNA [26]. A suitable beta value of 18 was calculated, and the TOMType was selected as “signed” to construct a coexpression matrix, which divided genes into 15 modules, including 14 valid modules and 1 “gray” module. The correlations between the modules and traits were calculated to select the modules for further analysis. Hub genes were selected using MM > 0.9 and GS > 0.8.

## Results

### Combined application of potassium and nitrate had a better alleviation effect on ammonium toxicity in rice roots

We found that the alleviation of ammonium toxicity in the roots of rice seedlings was much more enhanced when K^+^ and NO_3_^−^ were supplied together than when they were supplied separately (Fig 1A, B). Therefore, we speculated that there might be a synergistic mechanism by which K^+^ and NO_3_^−^ participate in ammonium toxicity alleviation. To confirm our hypothesis, we first demonstrated that neither Na^+^ nor Cl^−^ had any effects on ammonium toxicity and its alleviation (Fig 1B-D). Combined application of K^+^ and NO_3_^−^ had significantly stronger alleviation effects on root length, root surface area and root number than individual application (Fig 2). Under NH_4_^+^ toxicity treatment, the root cap was enlarged and grew abnormally, which was only restored by KNO_3_ (Fig 3B). Both cell production and cell expansion contributed to root growth. Under NH_4_^+^ toxicity treatment, root meristem size was reduced compared to the control and could be partially restored by KCl or NaNO_3_, while the meristem zone under the KNO_3_ alleviation treatment was longer than that in the control (Fig 3A, C), which indicated that KNO_3_ had a significant effect on improving cell division. Cells in the elongation zones of the control and KNO_3_ alleviation treatment were similar and were clearly elongated and rectangular, while cells in root elongation zones under the NH_4_^+^ toxicity and KCl or NaNO_3_ alleviation treatments were elliptical and irregular (Fig 3A). These results indicated that KNO_3_ could promote cell division and restore cell elongation better in the roots of rice seedlings under ammonium toxicity.

**Fig 1.**
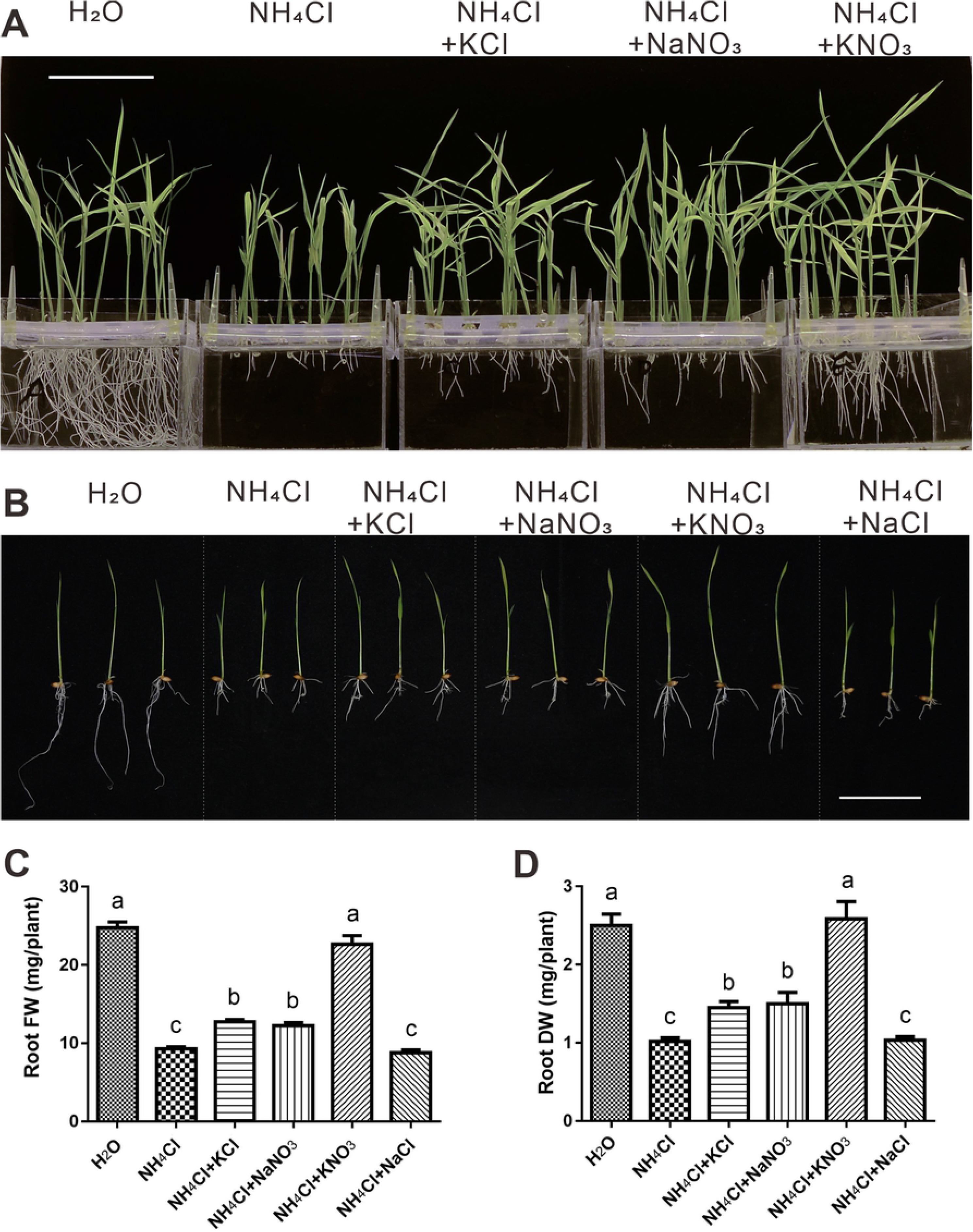
KNO_3_ had a better alleviation effect on NH_4_^+^ toxicity in rice roots than KCl or NaNO_3_. Phenotype of rice seedlings under different growth conditions for 10 days and 7 days (B). Fresh weight (FW) (C) and dry weight (DW) (D) of roots of 7-day-old seedlings. Scale bars = 5 cm. The data are the means ± SDs; t-tests were used to identify significant differences, and different letters represent significant differences among different treatments (p < 0.05).

**Fig 2.**
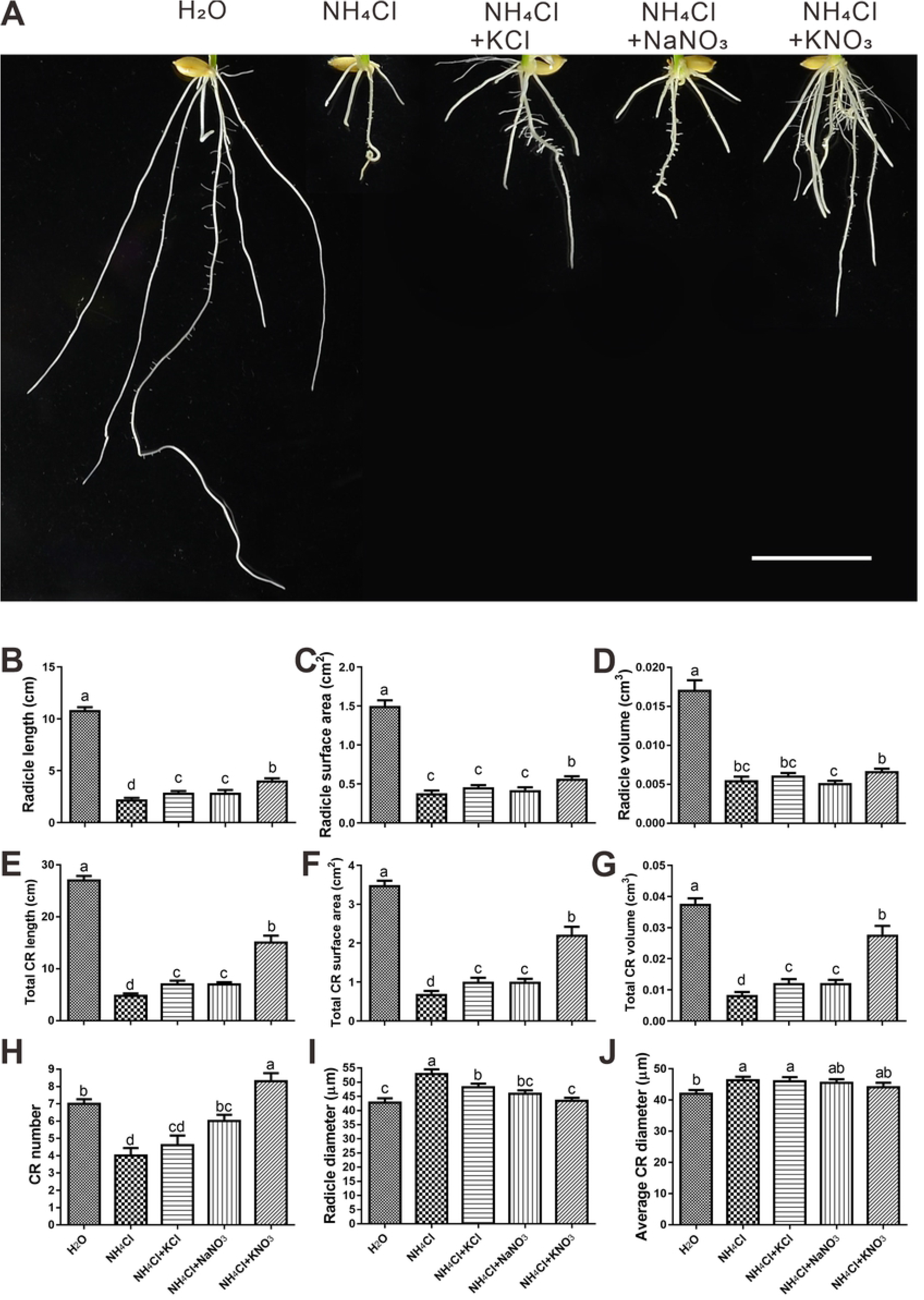
Root architecture statistics indicated significantly stronger alleviation effects when K^+^ and NO_3_^−^ were applied together. (A) Phenotypes of roots grown for 7 days. The length (B), surface area (C), volume (D) and diameter (I) of radicles were measured. The total length (E), surface area (F), volume (G) and average diameter (J) of the crown root (CR) were measured, and the number (H) of CRs was counted. Scale bar = 2 cm. The data are the means ± SDs; t-tests were used to identify significant differences, and different letters represent significant differences among different treatments (p < 0.05).

**Fig 3.**
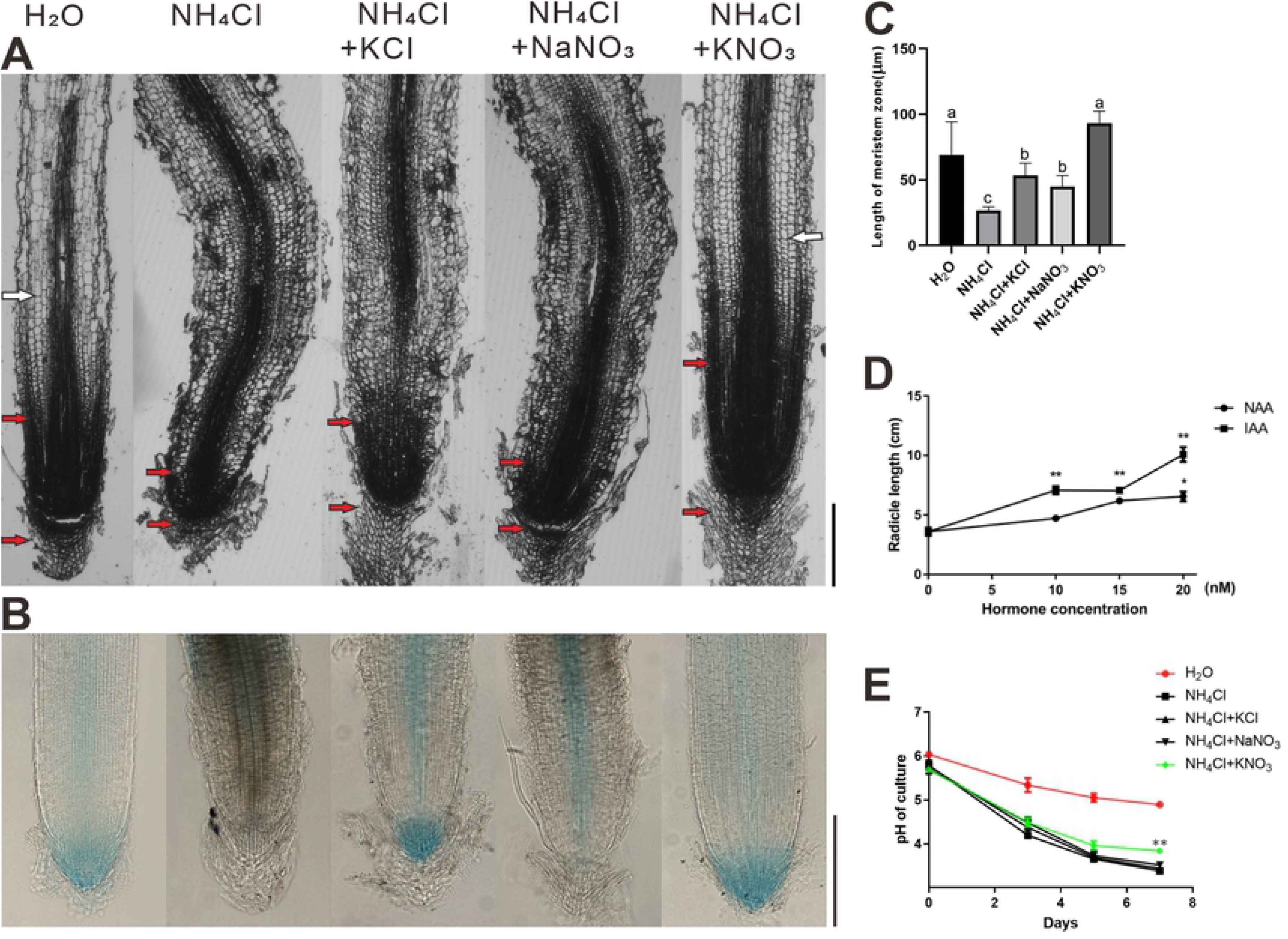
Only KNO_3_ could restore the cell phenotypes and auxin distributions of root tips and the rhizosphere acidification. Paraffin sectioning (A) and GUS staining (B) were performed on 7-day-old seedling radicles. The boundary of the meristem zone and elongation zone is marked by white arrowheads (A). The meristem zone (MZ) is marked by red arrowheads (A), and MZ lengths were measured (C). The roots shown in these photos were the most representative samples selected from 10 seedlings in each treatment. Rice seedlings were grown for 7 days under NH_4_^+^ toxicity treatment with different concentrations of IAA or NAA, and then, radicle lengths were measured (D). The pH of the culture solutions of different treatment was measured every 2 days from the 3^rd^ day after seed germination (E). Scale bars = 50 μm. The data are the mean ± SDs; t-tests were used to identify significant differences; different letters represent significant differences among different treatments (p < 0.05) for (C); “*” (p < 0.05) and “**” (p < 0.01) represent a significant differences compared to the control (0 nM hormone concentration) (for D), or between each alleviation treatment and NH_4_^+^ toxicity treatment (for E).

Under ammonium toxicity, the auxin signal in the rice seedling root tips was markedly weakened compared to that in the control (Fig 3B). KCl restored the distribution of the auxin signal in the root cap and significantly enhanced the auxin signal in the middle column of the root, while NaNO_3_ restored that in only the middle column of the root (Fig 3B). In the case of KNO_3_ alleviation, the distribution of the auxin signal was very similar to that in the control (Fig 3B), indicating that KNO_3_ had a better alleviation effect on auxin distribution in rice seedling roots. Addition of indole-3-acetic acid (IAA) or naphthaleneacetic acid (NAA) in the ammonium toxicity treatment increased the lengths of the rice radicles, which further indicated that the alleviation effect of KNO_3_ on root length may be attributed to auxin distribution recovery (Fig 3D). Rhizosphere acidification caused by ammonium toxicity could be alleviated by only KNO_3_ and not by KCl or NaNO_3_ (Fig 3E). Therefore, in terms of root phenotype, root biomass, root tip cell morphology, auxin distribution and rhizosphere acidification level, we demonstrated that the synergistic effect of K^+^ and NO_3_^−^ enhanced the alleviation of ammonium toxicity in rice seedling roots.

### The activities of ammonium assimilation metabolisms and the tissue ammonium levels in roots showed a same trend of differences between different treatments

To analyze how K^+^ and NO_3_^−^ alleviated rice NH_4_^+^ toxicity, we measured the NH_4_^+^ content in roots under different treatments. The NH_4_^+^ content under the ammonium toxicity treatment was the highest among all treatments (Fig 4A). Both KCl and KNO_3_ reduced the NH_4_^+^ content to a level comparable to that of the control, while the alleviation effect of NaNO_3_ was weaker (Fig 4A). These results suggested that both potassium and nitrate could alleviate root ammonium toxicity by reducing the net influx of ammonium. Furthermore, we compared the effects of K^+^ and NO_3_^−^ on root NH_4_^+^ influx and efflux using ^15^N-labeled NH_4_Cl. K^+^ was able to reduce NH_4_^+^ influx and efflux, while NO_3_^−^ was only able to reduce NH_4_^+^ efflux (Fig 4B, C). However, consistent with the NH_4_^+^ content, the impacts of KCl and KNO_3_ on root NH_4_^+^ influx and efflux were largely the same, suggesting that the differences in ammonium toxicity alleviation between KCl and KNO_3_ could not be attributed to the regulation of root NH_4_^+^ content. In addition, the GS, GOGAT enzyme activities and protein levels in the roots showed similar trends of differences among different treatment groups (Fig 4D-F).

**Fig 4.**
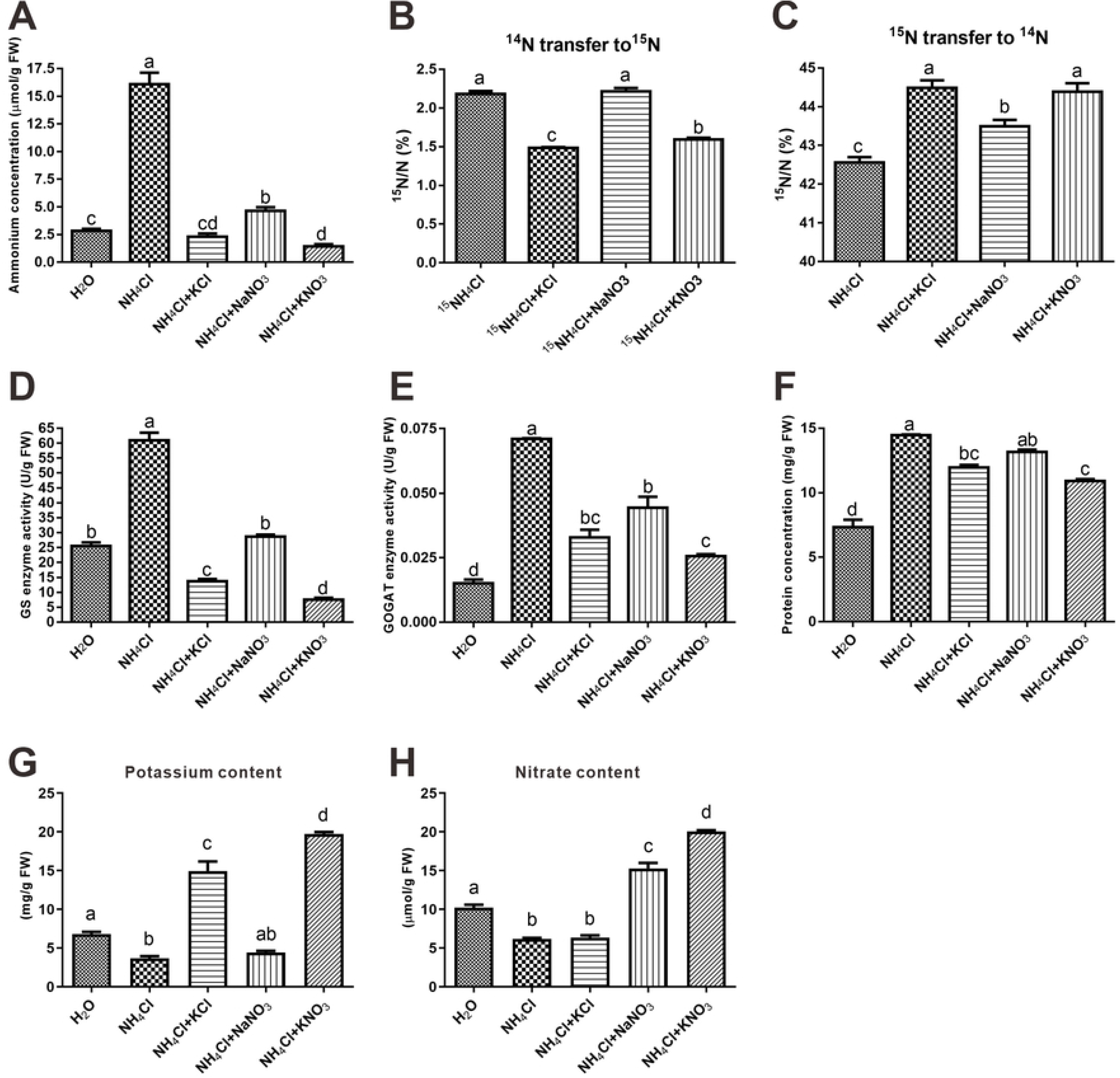
The activities of ammonium assimilation metabolisms and the tissue ammonium levels in roots showed a same trend of differences between different treatments. Rice seedlings were grown for 7 days under different treatments, and then, the roots were used to measure tissue ammonium (A), potassium (G), nitrate (H) and protein (F) levels, as well as to determine GS (D) and GOGAT (E) enzyme activities. For the ^15^NH_4_^+^ transport assay, rice seedlings were grown for 7 days in ^14^NH Cl (for B) or ^15^NH_4_Cl (for C) solution and then transferred to solutions containing ^15^NH_4_Cl (for B) or ^14^NH_4_Cl (for C) with the 3 alleviation treatments for 1 hour (the detailed experimental information is provided in the Materials and Methods) before the root ^15^N content was measured. The data are the means ± SDs; t-tests were used to identify significant differences, and different letters represent significant differences among different treatments (p < 0.05).

We further measured the K^+^ and NO_3_^−^ levels in roots under different treatments. The coexistence of K^+^ and NO_3_^−^ in the culture medium facilitated the absorption of both ions in rice roots (Fig 4G, H). However, the correlation analysis indicated that there was no correlation between K^+^ or NO_3_^−^ content and root growth (Fig 5).

**Fig 5.**
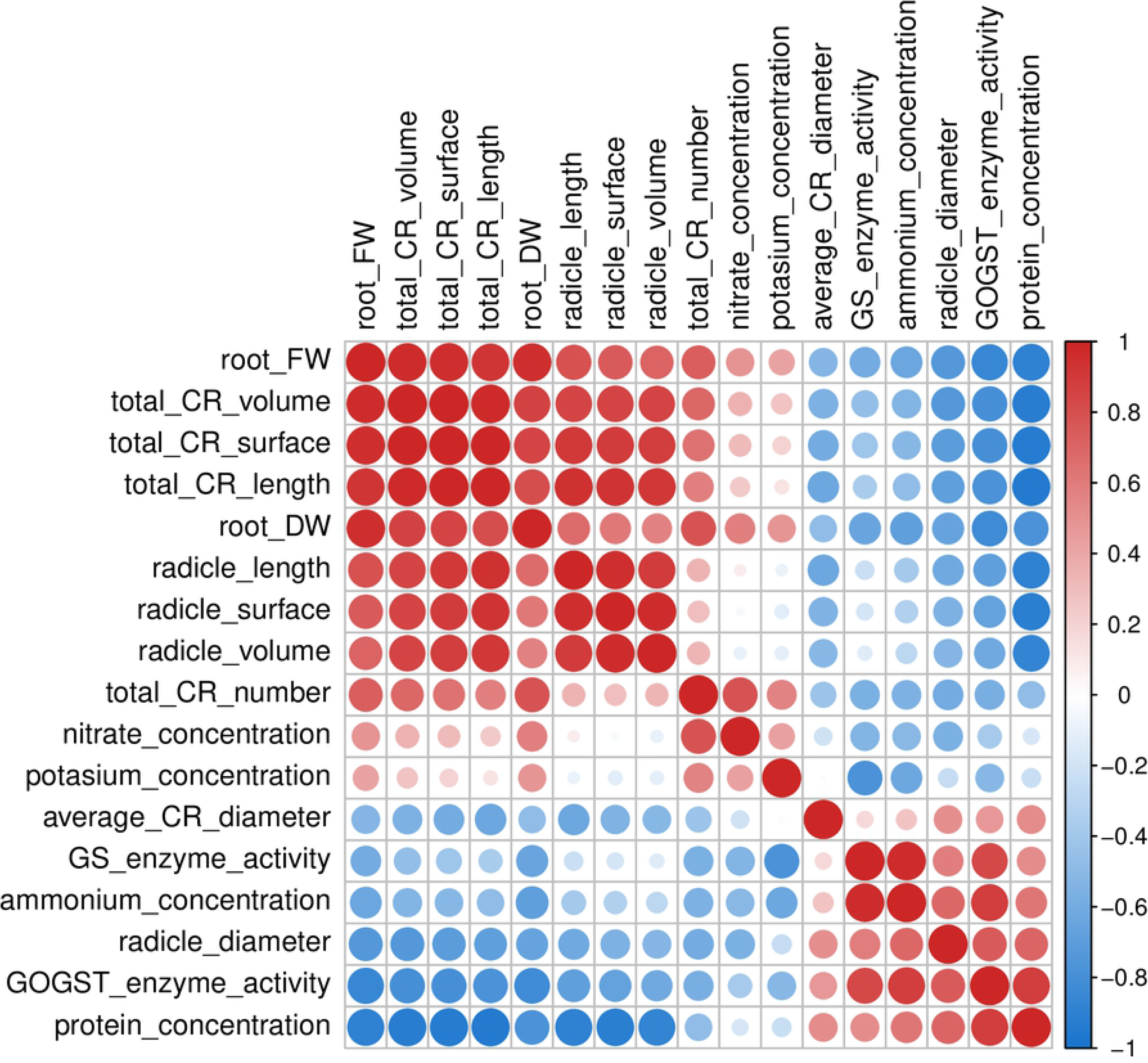
Correlation analysis revealed the correlations among phenotypic traits and physiological and biochemical data in rice roots. The color of each cell at the row-column intersections indicates the correlation coefficient between different traits or data. The color scale on the right side shows the correlation coefficient from −1 (blue) to 1 (red).

### Transcriptome analysis roughly revealed some related systems involved in this synergism

Whole-transcriptome analysis revealed that the number of differentially expressed genes (DEGs; |log2FC| > 1 and adjusted p value < 0.05) in each treatment compared to the control were in inverse proportion to the root biomass in corresponding treatment (Fig 6A, 1C-D). From these DEGs, we randomly selected 24 genes that appeared in all 4 groups to detect their expression level by quantitative real-time polymerase chain reaction (qRT-PCR), and all 24 gene expression differences among the 5 treatments were consistent with the corresponding fragments per kilobase per million (FPKM) values generated from transcriptome analysis, which verifies the validity of the RNA-Seq analysis results (S2 Fig). We defined 4365 DEGs between the NH_4_Cl treatment and the control as toxicity-related genes (Fig 6A). To further investigate the unique role of KNO_3_ in the alleviation of NH_4_^+^ toxicity, we compared the number of genes among the toxicity-related genes whose expression returned to levels comparable to that in the control (|log2FC| < 1 and adjusted p value < 0.05) under different alleviation treatments. There were 1716, 1335 and 2492 genes that met these requirements in DEG groups of the KCl, NaNO_3_ and KNO_3_ treatments, respectively (Fig 6B). These genes were treated as alleviation-related genes. There were 812 genes whose expression could only be recovered under KNO_3_ alleviation conditions (Fig 6B), further suggesting that a synergism existed between K^+^ and NO_3_^−^.

**Fig 6.**
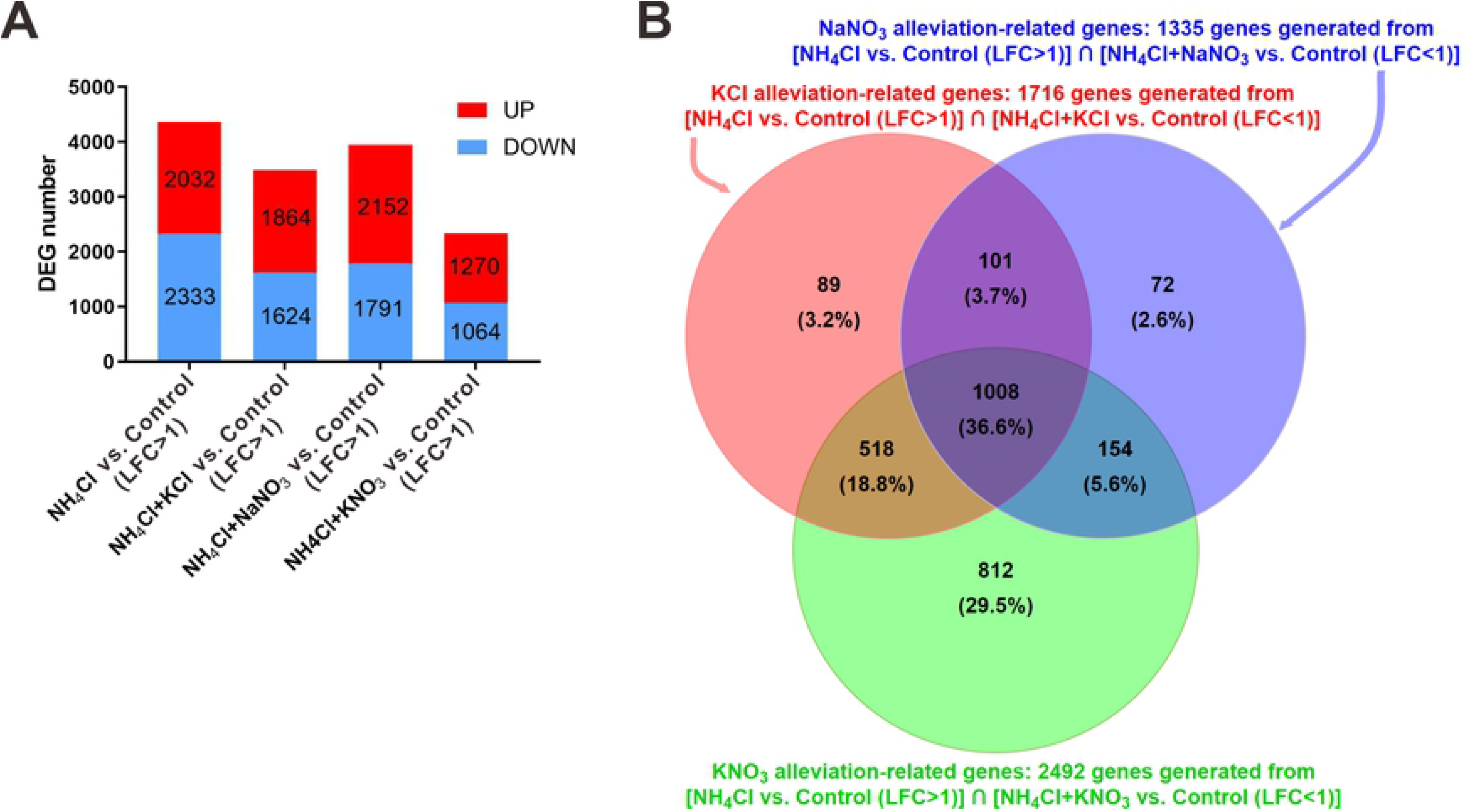
Study on this synergism using transcriptome analysis of the differentially expressed genes (DEGs). “LFC” represents |log2FC|. (A) Numbers of DEGs between different treatments and the control (H_2_O). (B) Venn diagram showing alleviation-related gene numbers in each alleviation treatment. The expression of these genes was differentially regulated by NH_4_^+^ toxicity and recovered to the control level by the 3 alleviation treatments.

To gain further insight into the possible processes involved in NH_4_^+^ toxicity and alleviation, we examined the distributions of toxicity-related genes and alleviation-related genes based on pathway enrichment (KEGG) analysis (S1A Fig) and gene ontology (GO) classification (S1B Fig). The results of KEGG and GO showed that the synergism of K^+^ and NO_3_^−^ generated new alleviation mechanisms via changes in gene expression related to stress responses, redox states and metabolism.

### Correlation network analysis by WGCNA allowed in-depth analysis of this synergism

To characterize the gene expression changes that occurred under different treatments, we clustered the expression patterns by the weighted gene correlation network analysis (WGCNA) method, leading to the identification of 15 distinct WGCNA modules (Fig 7A, C). Analysis of the module-trait relationships revealed that the ‘turquoise’ module was highly positively correlated with root biomass and all root traits except for root diameter (Fig 7C). The ‘turquoise’ module was also highly negatively correlated with GOGAT enzyme activity and protein concentration (Fig 7C). Analysis of trait correlation revealed that GOGAT enzyme activity and protein concentration were highly negatively correlated with all root traits except for root diameter (Fig 5). Therefore, the genes of the ‘turquoise’ module were considered to play an important role in root growth. The ‘yellow’ module was highly positively correlated with GOGAT enzyme activity, protein concentration and ammonium concentration (Fig 7C). The genes of the ‘yellow’ module might be negatively associated with root growth.

**Fig 7.**
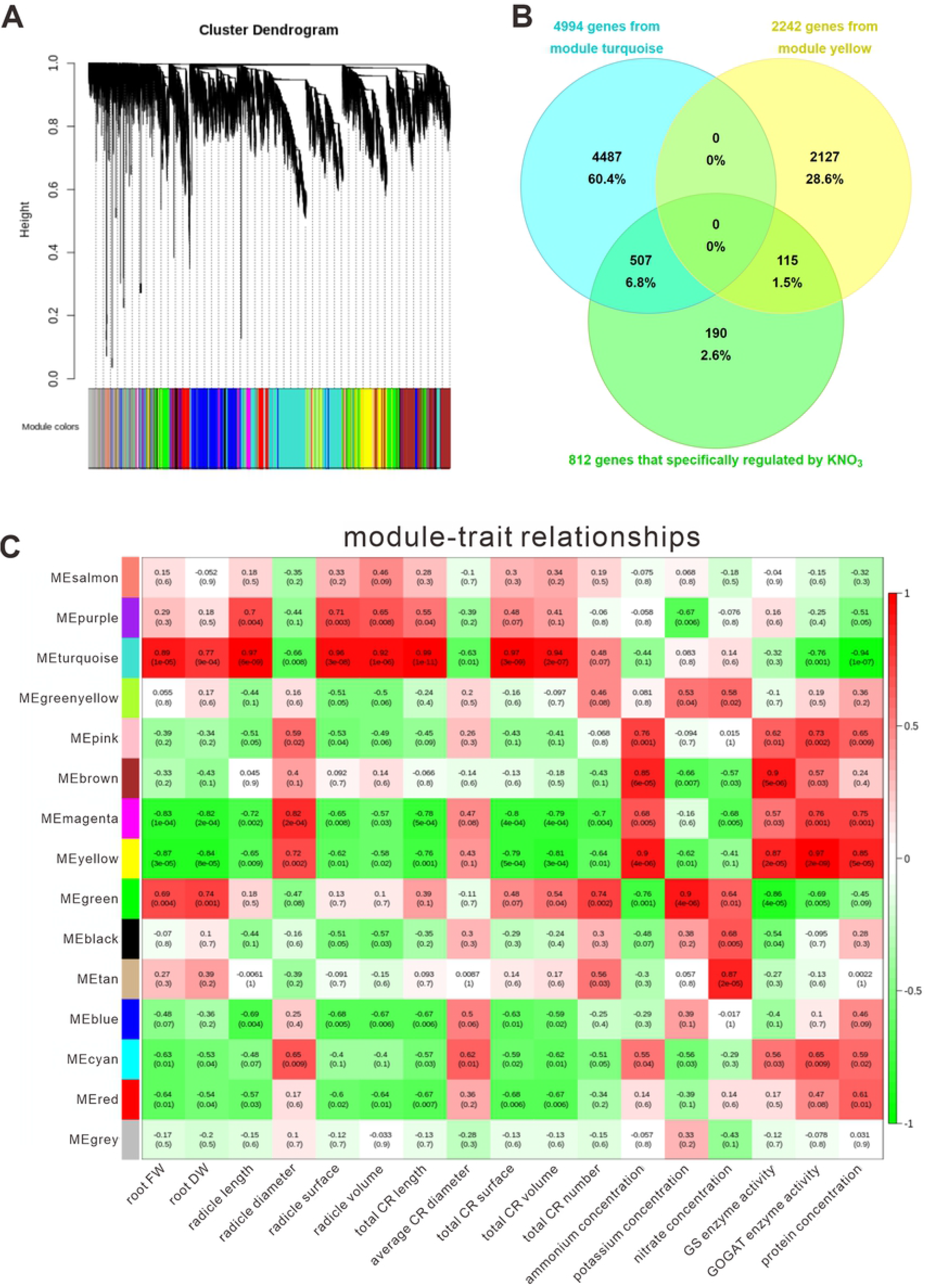
Study on this synergism using WGCNA. (A) Hierarchical cluster tree showing coexpression modules identified by WGCNA. Each leaf in the tree represents one gene. The major tree branch comprises 15 modules labeled by different colors. (B) Venn diagram showing the interactions of genes in root-trait coupled modules and DEGs whose expression was recovered only by KNO_3_ under NH_4_^+^ toxicity. (C) Module-trait association. Each row corresponds to a module. Each column corresponds to a trait. The color of each cell at the row-column intersection indicates the correlation coefficient from −1 (green) to 1 (red).

There were 507 genes in the ‘turquoise’ module and 115 genes in the ‘yellow’ module whose expression was differentially regulated by NH_4_^+^ toxicity and specifically recovered by KNO_3_ (Fig 7B). These genes whose expression was highly correlated with traits might be responsible for the specific synergistic alleviation effect on root growth. We used module membership (MM) > 0.9 and gene significance (GS) > 0.8 as parameters to identify hub genes among the 507 genes and found 62 hub genes (S1 Table). The coexpression network was visualized by Cytoscape with WGCNA edge weight > 0.35 (S1C Fig). Several genes involved in cell wall formation were identified in the network, such as genes associated with cellulose synthesis (OsCesA4, OsCesA7, OsCesA9), lignin catabolic process (Os01g0850700), cell wall modification (OsEXPB5) and cell wall proteins (OsFLA11, OsFLA27) (S1 Table). These results suggested that under ammonium toxicity conditions, KNO_3_ could specifically restore the expression of genes related to cell wall formation and then alleviate the effects of NH_4_^+^ toxicity on cell morphology and root growth by restoring cell wall formation.

### Construction of the cell wall regulation network related to this synergism

Based on these findings, we re-examined the expression of genes associated with root cell formation. Cellulose is an important component of the cell wall. Analysis of the transcription regulation network for cellulose synthesis revealed that OsNAC29 and OsNAC62 directly regulate OsMYB61, which in turn activates the expression of cellulose synthase genes (CESAs), such as OsCESA4, OsCESA7 and OsCESA9 [27]. The expression of OsMYB61 and OsCESAs is negatively regulated by OsIIP4, which does not bind their promoters and interacts with OsNAC29 [28]. Our result showed that the expression of these genes was repressed under NH_4_Cl treatment, and KNO_3_ resulted in a better recovery of the expression of these genes (Fig 8). Furthermore, similar results were observed with OsCSLF6 and OsIRX10 (Fig 8), which are in the hemicellulose synthesis pathway. Thus, the cellulose and hemicellulose synthesis pathways mentioned here might be a regulatory mechanism induced by the synergism.

**Fig 8.**
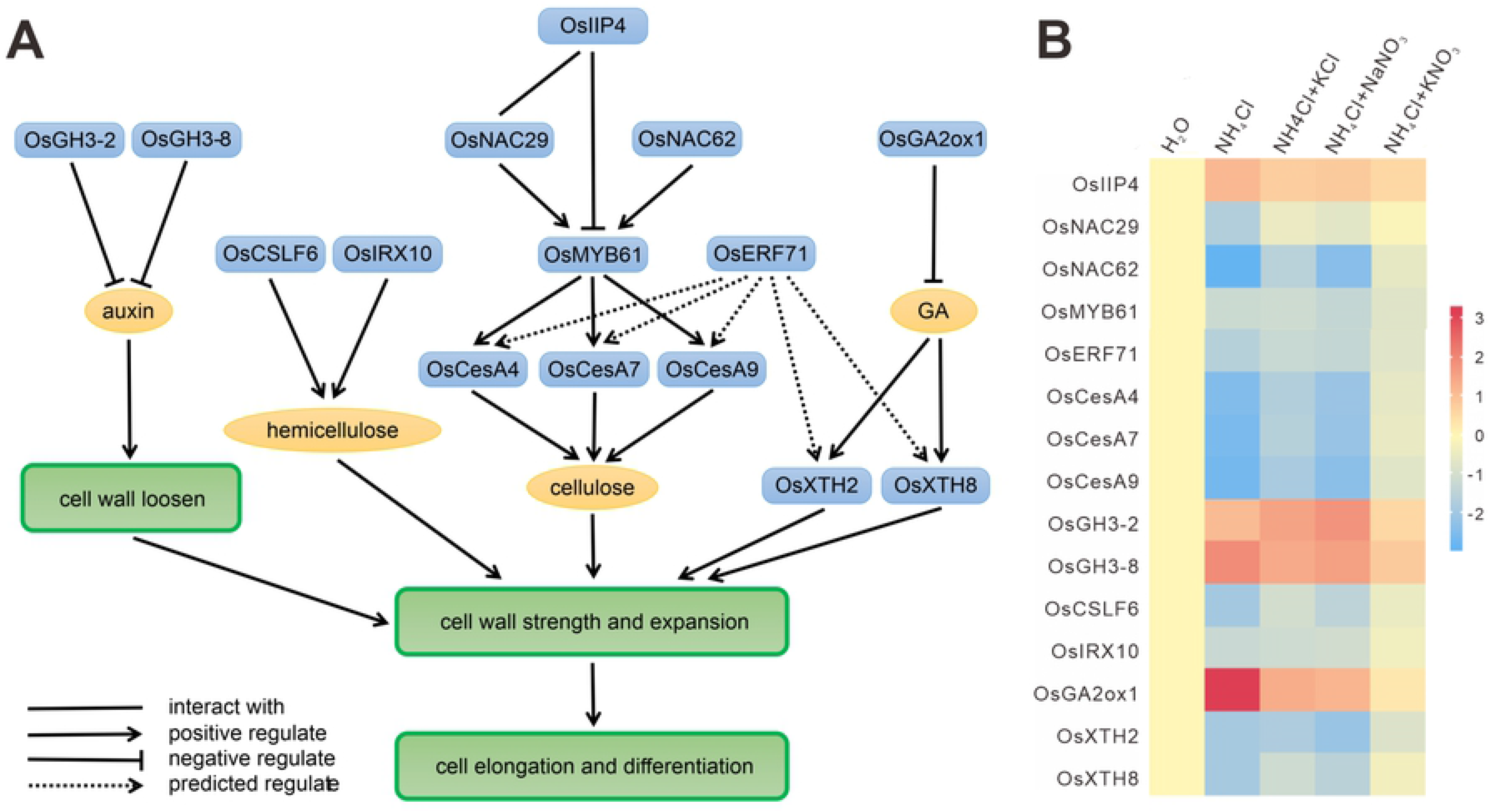
Construction of a cell wall regulation network related to this synergism. (A) A network involved in cell wall formation was constructed according to the literature (line) and predicted transcriptional regulation (dashed line). Blue boxes indicate genes or proteins; yellow ovals indicate chemicals; green boxes indicate biological processes. (A) Expression patterns of genes in this network under different treatments.

Xyloglucan endotransglucosylases/hydrolases (XTHs) play an important role in the construction of the primary cell wall by mediating the cleavage and rejoining of β-(1-4)-xyloglucans. There were two XTH genes (OsXTH8 and OsXTH2) among the 507 genes. OsXTH8 is upregulated by gibberellin (GA) and highly expressed in vascular bundles of the leaf sheath and young roots where cells are actively undergoing elongation and differentiation [29]. OsXTH2 is also upregulated by GA [30]. GA can be converted to nonbiologically active GA in plants by 2β-hydroxylation, a reaction catalyzed by GA 2-oxidase. OsGA2ox1 encodes a GA 2-osidase, which is functional in vitro and in vivo [31]. Our result showed that KNO_3_ could result in better recovery of the expression levels of OsXTH8, OsXTH2 and OsGA2ox1 under NH_4_Cl toxicity (Fig 8). This result suggested that the GA metabolic pathway mentioned here was disrupted by ammonium toxicity, leading to abnormal root cell elongation, which could be effectively recovered by the synergistic effects of KNO_3_.

Auxin is known to stimulate cell elongation by increasing cell wall extensibility. The OsGH3 protein controls auxin homeostasis by catalyzing the conjugation of IAA to amino acids. Overexpression of OsGH3-2 reduces the free-IAA level [32], and downregulation of OsGH3-8 shows auxin overproduction phenotypes [33]. Again, our result showed that the expression levels of OsGH3-2 and OsGH3-8 were upregulated by NH_4_Cl treatment, and high expression was maintained under the KCl and NaNO_3_ treatments (Fig 8). Only the KNO_3_ treatment restored the expression of these two genes to the control level, which was consistent with the auxin distribution result indicated by DR5:GUS (Fig 3B). Thus, the synergistic effects could also regulate the auxin pathways to result in enhanced beneficial effects on rice roots under NH_4_^+^ treatment.

## Discussion

### Potassium and nitrate had a synergistic effect on ammonium toxicity alleviation in rice roots

It is widely known that supplementation with nitrate or potassium alleviates ammonium toxicity in plants [1, 4, 16]. We found that the addition of KNO_3_ resulted in significantly enhanced promotion of root growth compared to that of KCl or NaNO_3_. In terms of root tip cell morphology, auxin distribution, and pH changes in the rhizosphere, the addition of KNO_3_ produced some alleviation effects that could not be produced by K^+^ or NO_3_^−^ alone, suggesting the existence of a synergistic effect of K^+^ and NO_3_^−^ on NH_4_^+^ toxicity alleviation.

Qing Li’s study concluded that ammonium inhibited root cell elongation rather than proliferation in plants [3]. Ying Liu’s study found that the inhibitory effect of ammonium on main-root elongation was mainly achieved by reducing the length of the meristem and elongation zones, and root cell elongation was also inhibited [5]. Our results showed that ammonium toxicity caused irregular cell morphology, impaired cell elongation, and loss of auxin signaling. KNO_3_ treatment effectively restored root cell morphology, resulting in a regular cell arrangement and a recovered auxin distribution that was almost the same as that of the control.

### This synergism could specifically regulate gene expression associated with root growth and ammonium assimilation

NH_4_^+^ causes changes in gene expression in plant roots, which in turn leads to a series of changes in root growth. K^+^ and NO_3_^−^ restored the expression of some of these genes to levels comparable to those in the control. Therefore, the restoration of the expression of these genes represents an intrinsic mechanism of ammonium toxicity alleviation by these two ions. We identified 812 genes whose expression was specifically restored by KNO_3_ but not by KCl or NaNO_3_, suggesting that the synergism between K^+^ and NO_3_^−^ activates specific transcriptional regulation to alleviate ammonium toxicity in roots.

Coexpression network analysis can be used for the investigation of thousands of genes with identical expression patterns. It is an important and excellent method for gene interaction analysis and interpretation of molecular mechanisms [34]. We identified a “turquoise” module that was highly positively correlated with almost all root traits (except for root diameter) and a “yellow” module that was highly positively correlated with tissue ammonium content and ammonium assimilation. The correlation analysis of the traits showed a negative correlation between most root traits and the ammonium assimilation. Therefore, we hypothesized that gene expression in these two modules is positively and negatively correlated with root growth. Among the 812 alleviation-related genes whose expression was specifically regulated by KNO_3_, 507 genes belonged to the ‘turquoise’ module and 115 genes belonged to the ‘yellow’ module. So the specific synergistic alleviation effect of KNO_3_ on root ammonium toxicity might be mainly achieved by regulation of the expression of these genes. No significant enrichment of GO terms was found in the 115 genes. The 507 genes were predominantly enriched in GO terms such as oxidation reduction, carbohydrate metabolic process, lignin catabolic process and response to oxidation stress. Furthermore, we identified 62 hub genes from the 507 genes. Mapman analysis [35] of these hub genes showed that they were enriched in three categories: genes associated with cell wall formation, various enzymes and transporters. Some genes were significantly associated with root growth traits and cell wall formation, so changes in their expression could result in different morphologies of root tip cells under ammonium toxicity or different alleviation conditions.

### This synergism alleviates root ammonium toxicity by regulating root cell wall formation

With changes in external nutrients, the composition of the cell wall determines the morphology of root cells and tissues, further influencing plant growth [36]. Although there is growing evidence that reprogramming of cell wall genes is critical for plant adaptation to nutritional status, an understanding of the molecular mechanisms that control these changes is now emerging. In crops, nitrogen status affects the mechanical strength and disease resistance of stems, which are regulated by cell wall organization and strength, suggesting that cell wall structure is regulated by nitrogen status [36, 37]. The expression of genes associated with lignin and cellulose synthesis is significantly downregulated when rice is grown in a high-nitrogen environment [37, 38]. Consistent with this phenomenon, a high-nitrogen growth environment causes a significant decrease in cellulose and lignin in the root cell wall, while nitrogen deficiency causes an increase in cellulose in roots [39].

Based on DEG analysis and WGCNA, we revealed a transcriptional regulatory network of genes associated with rice root cell wall formation that are affected by nutritional status (Fig 4, S2 Table). Our results showed that the cell wall formation in rice roots was related not only to nitrogen status but also to interactions between different nutrients. Separate addition of nitrate and potassium into the ammonium growth environment could not effectively alleviate the damage to cell wall formation, while the mixture of nitrate and potassium could effectively restore the expression of genes related to cell wall synthesis, thus restoring cell wall formation. Moreover, these processes also involve hormonal pathways such as the auxin and GA pathways. These results suggest that the effects of nitrogen on cell wall formation involve not only the morphological impact of nitrogen itself (ammonium or nitrate) but also the synergism between nitrogen and other nutrients (potassium). This is also consistent with the fact that, in agricultural practices, mixed fertilizer application results in better crop growth performance. An in-depth study of these mechanisms will reveal the link between environmental nutrition and crop growth and provide a theoretical basis for resolving the conflict between high yield and poor disease resistance in crops by improving fertilizer utilization efficiency.

## Acknowledgements

This work was supported by the National Key Research and Development Program of China [grant number 2016YFD0100400] and the National Special Key Project for Transgenic Breeding [grant number 2016ZX08001001].

